# Neural Synchronization as a Function of Engagement with the Narrative

**DOI:** 10.1101/2023.01.01.522416

**Authors:** Tal Ohad, Yaara Yeshurun

**Affiliations:** Sagol School of Neuroscience, Tel-Aviv University; School of Psychological Sciences, Tel-Aviv University

**Keywords:** DMN, Engagement with Narrative, fMRI, Valence, Inter Subject Representation Similarity Analysis, Inter Subject Functional Connectivity

## Abstract

We can all agree that a good story engages us, however, agreeing which story is good is far more debatable. In this study, we explored whether engagement with a narrative synchronizes listeners’ brain responses, by examining individual differences in engagement to the same story. To do so, we pre-registered and re-analyzed a previously collected dataset by Chang et al. (2021) of functional Magnetic Resonance Imaging (fMRI) scans of 25 participants who listened to a one-hour story and answered questionnaires. We assessed the degree of their overall engagement with the story and their engagement with the main characters. The questionnaires revealed individual differences in engagement with the story, as well as different valence towards specific characters. Neuroimaging data showed that the auditory cortex, the default mode network (DMN) and language regions were involved in processing the story. Increased engagement with the story was correlated with increased neural synchronization within regions in the DMN (especially the medial prefrontal cortex), as well as regions outside the DMN such as the dorso-lateral prefrontal cortex and the reward system. Interestingly, positively and negatively engaging characters elicited different patterns of neural synchronization. Finally, engagement increased functional connectivity within and between the DMN, the dorsal attention network and the control network. Taken together, these findings suggest that engagement with a narrative synchronizes listeners’ responses in regions involved in mentalizing, reward, working memory and attention. By examining individual differences in engagement, we revealed that these synchronization patterns are due to engagement, and not due to differences in the narrative’s content.

## 1. Introduction

Narratives are an essential component of human lives. We spend our days telling and hearing stories, to others and to ourselves. From children’s fables, through novels, to movies, humans are constantly inventing, sharing and evaluating stories. Stories can comfort, and can excite, they can divide, and they can unite. However, what might be enthralling to one individual, might be boring to another; what is endearing to some, can be annoying to others. In this study we set out to test the neural basis of individual differences in engagement with a narrative, examining whether engagement synchronizes listeners’ brain responses, and whether this synchronization relates to valence towards the story.

Engagement with a narrative includes various behavioral dimensions (Busselle & Bilandzic, 2009): *narrative understanding* (i.e., the cognitive aspect of comprehending the story), *attentional focus* (i.e., the unawareness of one’s concentration on the story), *emotional engagement* (i.e., feeling empathy or sympathy for the characters) and *narrative presence* (i.e., the sense of detaching from the actual presence whilst entering the narrative world). The latter dimension corresponds with a mental state described in previous studies regarding similar phenomena, namely transportation (Green & Brock, 2000) and flow (Sherry, 2004). Extant research exploring the neural correlates of narrative engagement mainly focused on the involvement of the Default Mode Network (DMN) in representing these dimensions.

Neuroimaging studies have thoroughly linked narrative processing to the DMN (Deniz et al., 2019; Nguyen et al., 2019; Simony et al., 2016; Yeshurun et al., 2021). For example, the DMN was involved in processing a narrative only when individuals understood the narrative (Ames et al., 2015; Honey et al., 2012; Ki et al., 2016; Lerner et al., 2011). Moreover, the DMN was more synchronized among participants who interpreted the story in the same (rather than different) manner (Bacha-Trams et al., 2017; Finn et al., 2018; Jääskeläinen et al., 2008; Nguyen et al., 2019; Yeshurun et al., 2017) and among participants who shared perspective regarding the narrative (Hakonen et al., 2022; Lahnakoski et al., 2014). The DMN’s role in attentional focus on a narrative was demonstrated in a study that found that the attended (and not the ignored) narrative synchronized participants’ DMN response (Regev et al., 2019), as well as in a study that documented higher neural synchronization among participants who paid attention to a narrative than participants who watched the same stimulus while counting down in steps of 7 (Ki et al., 2016). Additionally, it has been shown that identification with fictional characters may shape the response of regions within the DMN (Broom et al., 2021). These examples link behavioral dimensions of engagement to neural responses in regions of the DMN.

To explore how the brain response of listeners (or perceivers) changes as a function of their engagement with the narrative, previous research adopted two main paradigms: testing differences in the brain response to (i) more (vs. less) engaging narratives, and (ii) more (vs. less) engaging parts within the same narrative. Differences in the brain response were defined as differences in synchronization of the listeners’ (or perceivers’) neural activity. Studies testing for differences in the brain response found that engaging narratives synchronized the perceivers’ response within DMN regions. For example, neural activity within regions of the DMN was found to be more synchronized among participants who heard rhetorically powerful (compared to weak) speeches (Schmälzle et al., 2015), listened to personal (compared to non-personal) narratives (Grall et al., 2021) and were exposed to more (compared to less) effective stimuli regarding risky alcohol use (Imhof et al., 2017). Moreover, neural synchrony of participants who viewed a set of video clips was positively correlated to the amount of time other participants were willing to view them (Cohen et al., 2017).

The influence of engagement on neural synchronization was also studied from the perspective of dynamically changing degrees of engagement within the same narrative. For example, neural activity was found to be more synchronized among participants during arousing moments of the stimulus (Dmochowski et al., 2012). Furthermore, parts of a narrative that elicited negative valence were associated with increased synchronization in regions of the DMN (Nummenmaa et al., 2012, 2014). In a recent study by Song et al. (2021), a set of participants defined the engaging moments of two narratives, and another set of participants underwent an fMRI scan while listening to the same two narratives. Neural synchronization within regions of the DMN was higher during more engaging moments of the narratives.

Engagement with a narrative has also been associated with enjoyment (Green et al., 2004). The measurement of participants’ transportation into a story was highly correlated with the enjoyment that was experienced during attending to it (Busselle & Bilandzic, 2009; Green et al., 2004). These assosciations give credence to the view that while enjoyment is not a prerequisite for engagement, it may be a significant factor. Yet, the involvement of the reward system, which is activated by stimuli that elicit enjoyment (Klasen et al., 2012; Wabnegger et al., 2021), has not been studied in the context of narrative esngagement. More on this front, it has been shown that engagement can be generated via both positive emotions (Jin-Ae et al., 2020) and negative emotions (Landreville & LaMarre, 2011). However, previous studies did not test whether the same neural mechanisms support both positive and negative emotional engagement.

## 2. Hypotheses and Goals

The aforementioned research suggests that more engaging narratives enhance neural synchronization in regions within the DMN. While these studies shed light on the brain correlates of narrative engagement, it is important to note that they compared the brain response to different stimuli content (e.g., different narratives or different parts of the same narrative), and thus differences in brain response may also emerge due to differences in the content (e.g., more versus less emotional content). In this project we suggest a complementary approach to study narrative engagement, in which we take advantage of individual differences in engagement with a fixed narrative, thereby avoiding content-dependent differences in the brain response.

In the present pre-registered study, we hypothesized that neural responses within regions of the DMN would be more synchronized among participants who were more engaged with the narrative. We further hypothesized that the neural responses within regions of the DMN and regions associated with reward and valence would be more synchronized during an appearance of a specific character among participants who were more engaged with the narrative, regardless of the emotional valence that the character induced. Moreover, we explore whether functional connectivity within the DMN and between the DMN and other brain networks depends on participants’ engagement with the narrative. To address these questions and predictions, we used an fMRI dataset of participants who listened to a one-hour narrative and rated their engagement with it (Chang et al., 2021). We employed Inter-Subject Correlation (ISC; Hasson et al., 2004), Inter-Subject Representational Similarity Analysis (IS-RSA; Finn et al., 2020) and Inter-Subject Functional Connectivity combined with Representational Similarity Analysis (ISFC-RSA; Finn et al., 2020; Simony et al., 2016) to test for engagement-dependent brain response.

## 3. Methods

The dataset, hypotheses and plan for analysis in the present study were pre-registered on the Open Science Framework (OSF). This was done after data collection but prior to data analysis (https://archive.org/details/osf-registrations-keuja-v1).

### 3.1. Participants

The current pre-registered study re-analyzes a previously published fMRI dataset (Chang et al., 2021) consisting of fMRI scans of 25 right-handed participants (13 females, 12 males; age: *M* = 22.48, *SD* = 4.95).

### 3.2. Stimulus and Experimental Design

The narrative stimulus was a 56-minute recording of a professional actor (June Stein) reading a story by Christina Lazaridi, “The 21^st^ Year.” The narrative consisted of 45 segments, each lasting for 61–86 seconds (*M* = 70 sec, *SD* = 4.2 sec), that were separated by silent pauses of 4–6 seconds. It included six main characters (Clara, Gary, Sasha, Margaret, Steven and Alexander) that appeared throughout the story (Fig. 1). Following the scan, outside the scanner, participants answered a detailed questionnaire about the story they just heard.

**Fig. 1.**
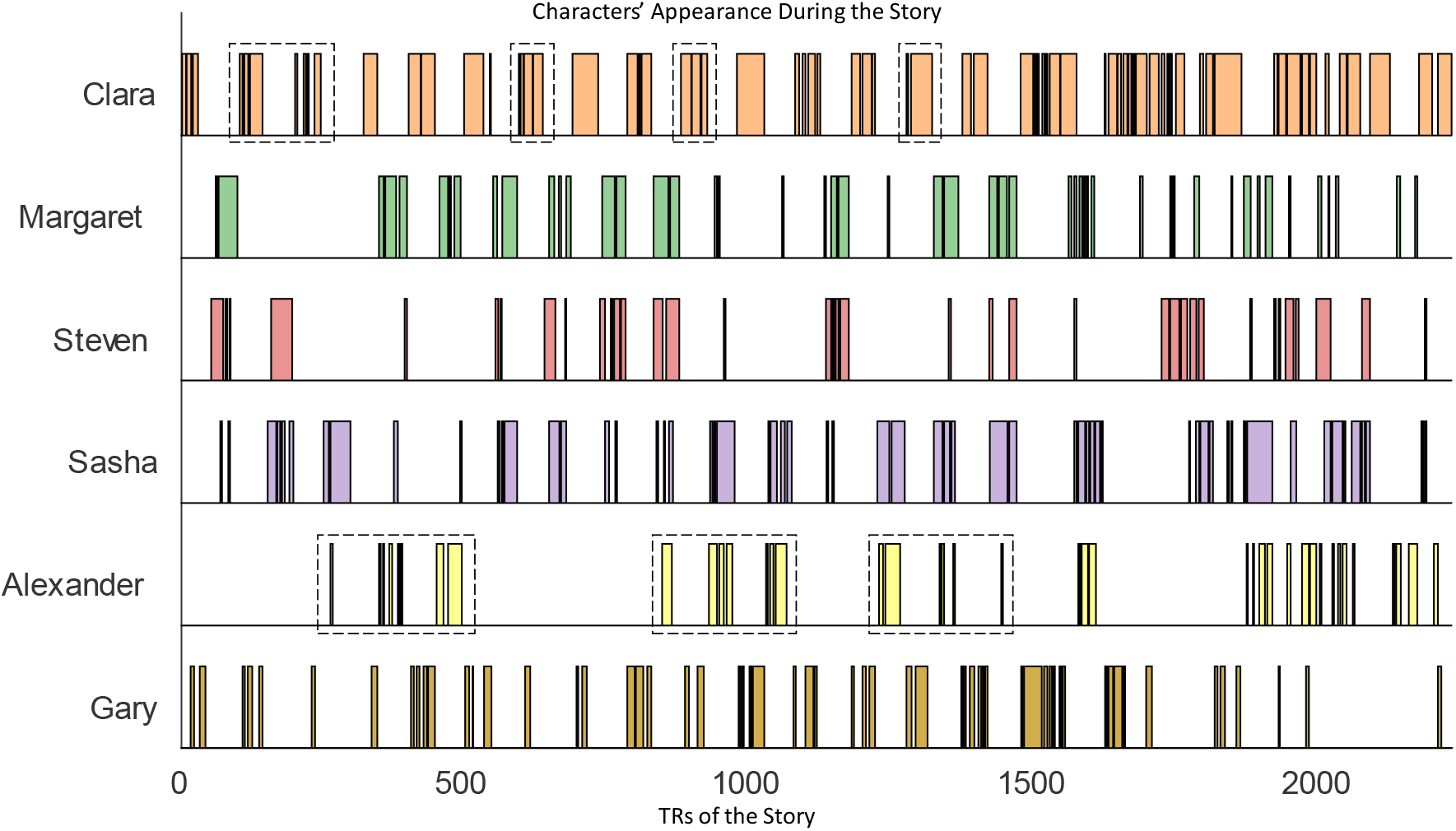
Appearance of every character along the timeline of the story in TRs. The dotted rectangles represent the TRs that were taken into account for analyzing the characters.

### 3.3. Post-Scan Questionnaire

The detailed post-scan questionnaire included questions regarding personal aspects that might be relevant to the individuals’ interpretation of the narrative, as well as questions regarding their memory, engagement and reactions to the narrative. In this study, to assess participants’ engagement with the narrative, we used the questions that evaluated participant’s overall engagement with the story (on a scale of 0 to 4), the degree of engagement (on a scale of 0 to 4) with each of the five main characters (Clara, Sasha, Margaret, Steven and Alexander; note that due to technical reasons we did not measure engagement with Gary), whether their engagement with these characters was positive, negative or neutral and the reasons for these engagement ratings.

To test for similarities and differences in positive and negative engagement-dependent neural responses, we chose one character that elicited positive engagement and one that elicited negative engagement (based on the behavioral data). To better assess the difference between participants’ responses to these two characters, we used text analysis on their open answers to the questions regarding the reasons for engagement ratings and engagement valence. This analysis was done by Linguistic Inquiry and Word Count (LIWC-22; Boyd et al., 2022), a validated platform for evaluating meaning of written content. The words constructing the text were assigned to various linguistic categories, based on previously designed lexicons. To accommodate participants’ variations in text length, the amount of words in each category was provided as a percentage. We calculated the percentage of words that fit under nine categories: affiliation (e.g., we, help), positive tone (e.g., good, well), prosocial behavior (e.g., care, thank), negative tone (e.g., bad, wrong), anxiety (e.g., worry, fear), anger (e.g., hate, frustrated), sadness (e.g., disappointed, cry), affect (e.g., new, love) and moralization (e.g., wrong, deserve). We then used dependent t-tests to assess the difference between the two characters in relation to these categories. The *p*-values of the nine t-tests were then FDR-corrected (Benjamini & Hochberg, 1995).

### 3.4. Preprocessing

MRI data were preprocessed using FSL 5.0 (fsl.fmrib.ox.ac.uk/), including the Brain Extraction Tool (Smith, 2002), slice time correction, motion correction, high-pass filtering (140-sec cutoff) and spatial smoothing (FWHM = 6 mm). All data were aligned to standard 3-mm Montreal Neurological Institute (MNI) space (MNI152). Only voxels covered by 90% of all participants’ image acquisition area were included for further analysis. This criterion yielded a total of 50,530 voxels. After preprocessing, the first 16 TRs (repetition time; TR = 1.5 seconds) were cropped to remove the time gap between scanning and narrative onset. To correct for the hemodynamic delay and align the fMRI data to the stimulus, we followed Chang et al.’s (2021) procedure and cropped three additional TRs.

### 3.5. Regions of Interest (ROIs)

Pre-registered ROIs were defined using Neurosynth meta-analysis of the term “DMN” (including bilateral temporal parietal junction or TPJ, precuneus, dorsomedial prefrontal cortex or dmPFC, ventromedial prefrontal cortex or vmPFC and bilateral temporal poles); the term “reward” (including bilateral nucleus accumbens, bilateral amygdala and vmPFC); and the term “valence” (including bilateral amygdala and vmPFC). As there was an overlap of at least two of these terms in vmPFC and bilateral amygdala, we generated four non-overlapping maps: 1) vmPFC; 2) bilateral amygdala; 3) the remaining non-overlapping regions of the reward system; and 4) the remaining non-overlapping regions of the DMN system.

Additionally, we performed an exploratory whole-brain analysis, in which cortical regions were allocated into 114 parcels following Yeo et al. (2015). This parcellation is based on a 17-network cortical parcellation estimated from the resting-state functional data of 1,000 adults (Yeo et al., 2011; Yeo et al., 2015). Subcortical regions were allocated into 8 parcels, corresponding to bilateral amygdala, hippocampus, thalamus and striatum, extracted from the subcortical nuclei masks of the Brainnetome atlas (Fan et al., 2016). The time courses of the voxels corresponding to each of these 122 parcels (114 cortical and 8 subcortical) were averaged to obtain a single, representative time course. The 18 functional networks were divided into nine groups: visual, somatomotor, dorsal attention, ventral attention, limbic, control, temporal parietal, default mode, and the sub cortex (<u>Yeo et al., 2011</u>; Yeo et al., 2015). Each parcel was labeled with one of the nine functional networks.

### 3.6. Data Analysis

All analyses were performed on pre-registered ROIs and on the 122 parcels (defined by Yeo et al., 2015, and Fan et al., 2016).

#### 3.6.1. Inter-Subject Correlation (ISC)

To test whether a certain voxel (or parcel) was involved in processing the narrative, we performed ISC analysis (Hasson et al., 2004). We extracted each participant’s time course within each voxel (or parcel) for each of the 45 segments of the narrative. We then correlated this time course with the averaged time course of the remaining participants. Next, we averaged these ISC values across participants and segments. This resulted in one ISC value for each voxel (or parcel) (Fig. 2A).

**Fig. 2.**
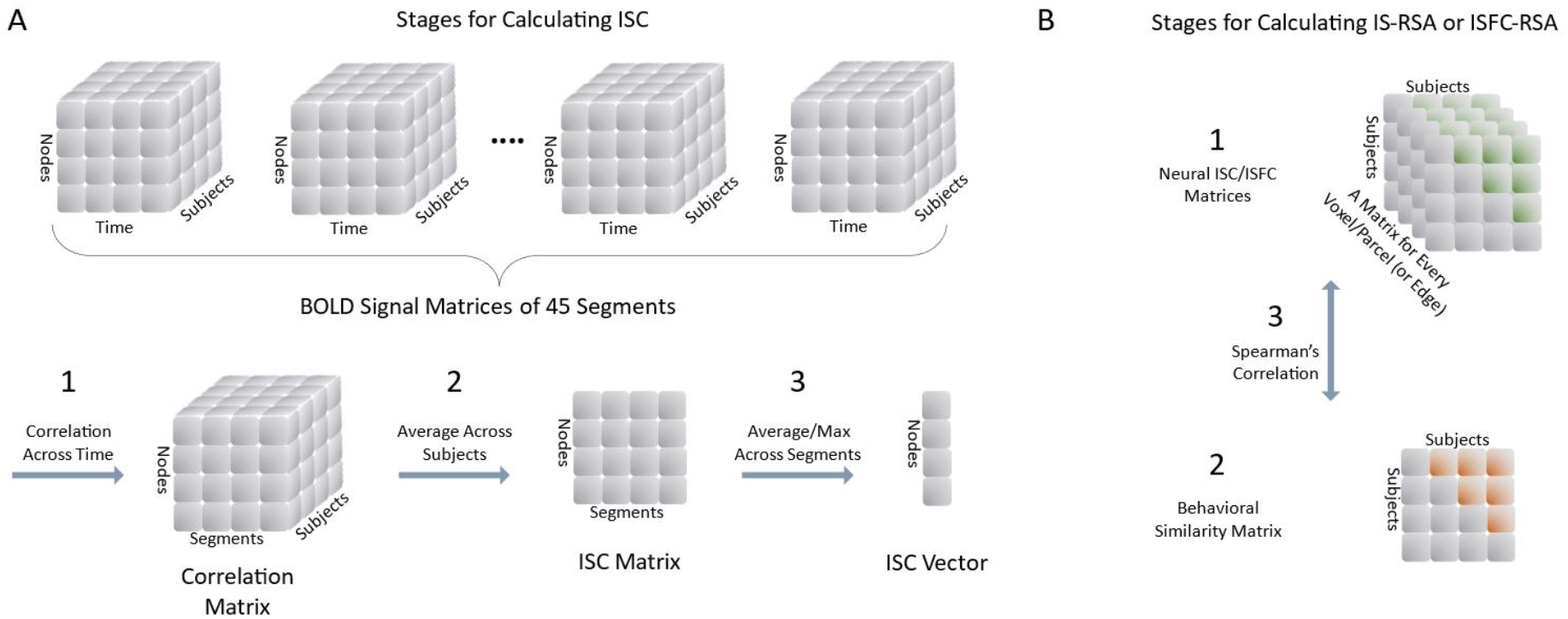
Methods. A) In order to calculate real ISC values, we followed these steps. 1: Within each voxel or parcel, within each segment, we correlated each subject’s time course with the averaged time course of the remaining subjects; 2: We averaged these correlation values across all subjects; 3: We averaged the ISC values across segments. For null values, on every iteration we phase-randomized the time course for every subject, within every voxel or parcel, for every segment, and followed steps 1-2. On step 3 we chose the segment with the maximal ISC value to constitute the null distribution of ISC. B) In order to calculate IS-RSA (or ISFC-RSA) values, we followed these steps. 1: For every voxel or parcel (or edge), we calculated the correlation between every two subjects to create neural matrices; 2: For every two subjects, we found the averaged score of engagement to create the behavioral matrix; 3: We used Spearman’s correlation to find the similarity between the behavioral matrix and every neural matrix.

To test whether this value was significantly greater than chance, we generated a null distribution for each voxel or parcel by randomizing the phase of the signal in 1,000 iterations, and then performing the same ISC calculation on the shuffled signal (Simony et al., 2016). Within every iteration, among the 45 ISC scores (one for each segment), the maximal ISC score was included in the null distribution. We chose to take the maximal and not the averaged ISC score to obtain a more conservative estimation of the chance level. To estimate the *p*-value of a voxel’s (or parcel’s) real ISC value, we calculated the complementary cumulative distribution function of the null distribution, evaluated at the real ISC value (Fig. 2A). We selected only voxels (or parcels) that achieved a significant ISC score (*q* < 0.05, corrected for multiple tests using FDR).

#### 3.6.2. Inter-Subject Representational Similarity Analysis (IS-RSA)

To test whether higher engagement with the narrative and with specific characters was associated with more similar brain responses, we performed IS-RSA analysis (Fig. 2B; Finn et al., 2020).

#### IS-RSA for engagement with the story

Similarity in engagement intensity was calculated using the AnnaK model (Finn et al., 2020). According to this model, similarity between subjects is best represented by their average, as it assumes that similarity between subjects increases or decreases as one proceeds up or down the scale. Thus, for each pair of participants, we calculated their averaged engagement with the narrative. This resulted in a 25×25 behavioral similarity matrix. Similarity in the brain response for the whole narrative was calculated by computing subject-pairwise correlations of hemodynamic responses in each voxel (or parcel) across the whole story (2,227 TRs). The IS-RSA was then computed as the Spearman correlation between the behavioral similarity matrix and the voxels’ (or parcels’) brain activity similarity matrices (Fig. 2B).

##### 3.6.2.1. Testing significance

To test whether voxels’ (or parcels’) correlation value was significantly above chance, we combined two methods. First, in a similar procedure as was done for the ISC values, we generated a null distribution and calculated the *p*-value of the IS-RSA correlation and corrected these *p*-values using an FDR correction. Next, we wanted to verify that the obtained significant voxels (or parcels) were indeed involved in processing the narrative in at least some of the participants. To do so, we divided the 25 participants into three groups, based on their overall engagement ratings: the Low-Engaged group consisted of participants with an overall engagement score of 0–2 (N = 9), the Medium-Engaged group consisted of participants with an overall engagement score of 3 (N = 9), and the Highly Engaged group consisted of participants with an overall engagement score of 4 (N = 7). We then calculated the ISC for each of these three groups and selected voxels (or parcels) that passed the threshold in at least one group. The threshold was the 99^th^ percentile of the null distribution of the ISC (DMN – 0.1748, reward – 0.1488, amygdala – 0.1411, vmPFC – 0.1564, whole-brain parcellation – 0.1621).

##### 3.6.2.2. IS-RSA for engagement during characters’ appearance

We performed IS-RSA for engagement during the appearance of two of the main characters (Clara and Alexander). These two were chosen based on the valence of the engagement they induced among the participants (one positive – Clara, and one negative – Alexander), with a preference given to the largest variability in engagement ratings (see Fig. 4).

Similarity in engagement intensity was calculated using the AnnaK model (Finn et al., 2020). In two separate analyses (one per character), for each pair of participants, we calculated their averaged overall engagement, which resulted in one 25×25 behavioral similarity matrix. As to similarity in the brain response for a specific character, we first extracted the TRs in which a specific character appeared in the story without the other character and concatenated these TRs (542 TRs for Clara and 178 TRs for Alexander; see Fig. 1). To avoid any differences that will stem from differences in number of TRs, we chose 178 TRs from Clara’s TRs that resembled Alexander’s TRs the most – both in density and in quantity (see Fig. 1). Next, we calculated subject-pairwise correlations of the hemodynamic responses within each voxel (or parcel) across a specific character’s TRs. The IS-RSA was then computed as the Spearman correlation between the behavioral similarity matrix and the voxel’s (or parcel’s) brain activity similarity matrices (Fig. 2B).

##### 3.6.2.3. Testing significance

As in the IS-RSA analysis for engagement with the story, to test whether a voxel’s (or parcel’s) correlation value was significantly above chance, we combined two methods. First, in a similar procedure as was done for the ISC values, we generated a null distribution and determined the *p*-value of the IS-RSA correlation. Next, to verify that the voxel (or parcel) was indeed involved in processing the narrative in some of the participants, we selected voxels (or parcels) that passed the threshold in at least one of the three groups (Low-, Medium- and Highly Engaged). The threshold was the 99^th^ percentile of the null distribution of the ISC (for Clara: DMN – 0.1791, reward – 0.1437, amygdala – 0.1412, vmPFC – 0.1518, whole-brain parcellation – 0.1611; for Alexander: DMN – 0.1746, reward – 0.1441, amygdala – 0.1418, vmPFC – 0.1506, whole-brain parcellation – 0.1632).

#### 3.6.3. Inter-Subject Functional Connectivity RSA (ISFC-RSA)

To test whether higher narrative engagement was associated with stronger connectivity between regions that demonstrated engagement-dependent response, we used ISFC-RSA (Fig. 2B; Finn et al., 2020; Simony et al., 2016). To do so, we first defined regions of interest (ROIs) as the voxels that appeared to be significant in the IS-RSA analysis, clustered by the Neurosynth maps (voxels in the DMN map were assigned to the PCC, PFC and left and right TPJ), and as the 29 parcels that were revealed by the whole-brain IS-RSA analysis to have engagement-dependent response (supplemental Table 2). Next, we estimated the connectivity between these ROIs by computing Inter-Subject Functional Connectivity (ISFC; Simony et al., 2016). For each of the ROIs, for each participant, we extracted the time course and correlated it with the averaged time course of the remaining participants, in each of the ROIs. These correlation values underwent a Fisher’s r-to-z transformation, were averaged across participants and then inverse-transformed to produce averaged correlation values. This resulted in a square ISFC matrix of the functional connectivity among these ROIs, and it was asymmetric due to the directional nature of this procedure. However, functional connectivity is considered to be unidirectional. Therefore, the symmetry was achieved by averaging the upper and lower triangles (Simony et al., 2016).

Similarity in intensity of overall engagement was calculated as in the IS-RSA analysis, using the AnnaK model (Finn et al., 2020), and resulted in a 25×25 behavioral similarity matrix. A neural similarity matrix was generated for every edge between every two ROIs by computing correlations of hemodynamic responses between the two specific ROIs between every two participants (a pairwise version of the ISFC analysis). This resulted in an ISFC matrix for every edge (29 × 28 ÷ 2 = 406 edges). The ISFC-RSA values were then computed as the Spearman correlations between the behavioral matrix and the edges’ neural similarity matrices (Fig. 2B).

To test whether the edges’ correlations were significantly above chance, we combined two methods. First, as in the ISC analysis, we generated a null distribution and calculated the *p*-values of the ISFC-RSA values. Second, as in the IS-RSA analysis, out of the significant edges (after FDR correction) we selected only edges that passed a threshold that was the 99^th^ percentile of the null distribution of the ISFC (Neurosynth maps – 0.1364, parcellation – 0.1658) in at least one of the three groups: Low-Engaged, Medium-Engaged and Highly Engaged participants.

### 3.7. Network Visualization

In order to visualize the ISFC-RSA results, we used the open-source program Cytoscape (Shannon et al., 2003) and the plot_connectome function from the Nilearn library of Python (Abraham et al., 2014). Each node was represented by a circle, the diameter scaled with the IS-RSA value of that node (diameter = 300 × IS-RSA; diameter = 200 × IS-RSA in Python and Cytoscape, respectively). The thickness of lines in the diagram indicates the strength of edges of the network (line width = 25 × edge-strength; Fig. 7).

### 3.8. Surface Display

Projections onto a cortical surface for visualization were performed with Human Connectome Project Workbench software (Marcus et al., 2011).

### 3.9. Ethics Statement

Data were obtained from the OpenNeuro dataset : https://openneuro.org/datasets/ds002245. All participants provided written informed permission, as described by Chang et al. (2021).

### 3.10. Code Availability

The code disclosed in this work is accessible from the SOCON’s lab GitHub repository at https://github.com/YeYeLab/SOCON-lab.git.

## 4. Results

### 4.1. Behavioral Results

Post-scan ratings revealed a large variability in participants’ overall engagement with the narrative (overall engagement: *M* = 2.8, *SD* = 1.02) (Fig. 3A). To characterize the valence of the engagement with the characters, we used participants’ ratings on whether their engagement with each character was positive, negative or neutral. These ratings revealed that Clara, Margaret, Steven and Sasha engaged participants in a positive manner (respectively, 80%, 60%, 40% and 76% of participants indicated positive engagement), and engagement with Alexander was mainly negative (88% of participants indicated negative engagement; Fig. 3B).

**Fig. 3.**
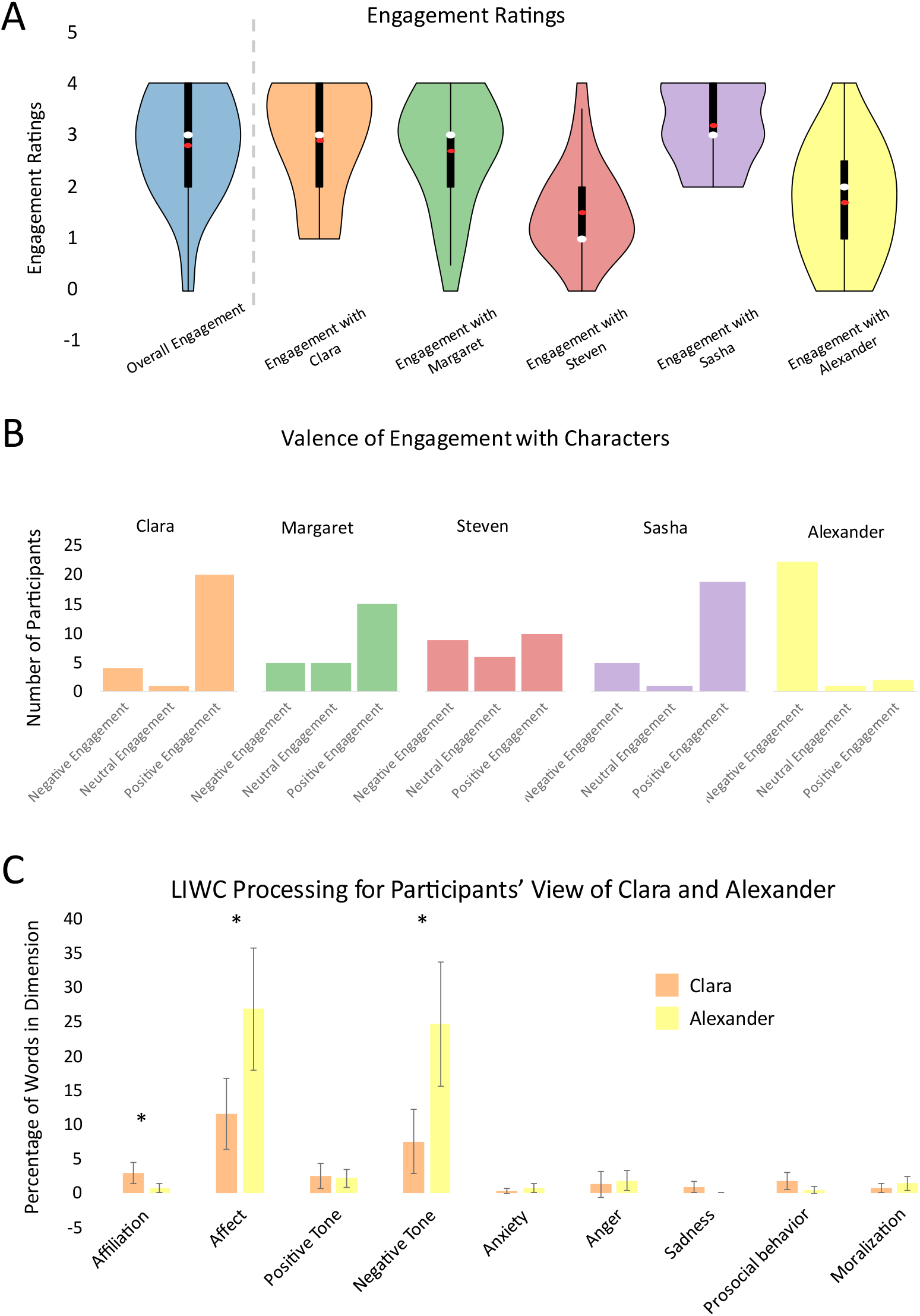
Behavioral results. A) Engagement ratings represented on a violin plot. The white dot is the median and the red dot is the mean. The bold line indicates the interquartile range. B) Histograms of the valence of engagement with every character. C) In the LIWC analysis for participants’ view of Clara and Alexander, the X axis indicates the categories, and the Y axis indicates the extent to which each category was represented in the participants’ answers. The error bars indicate the confidence interval with a confidence level of 95%.

For further analyses on how the valence of the engagement shapes the neural responses, we chose one character that induced positive engagement (Clara) and one that induced negative engagement (Alexander). To demonstrate what we mean by inducing positive and negative engagement, here are some examples from listeners’ responses regarding the valence of their engagement with Clara: “positive, she was relatable”; “positive, she was doing her best to save her marriage”; “positive because it made me want to know what happened next”. Some examples for listeners’ answers regarding the valence of their engagement with Alexander: “negative, he took advantage of both Margaret and Sasha”; “negative, he seemed selfish and a bit sadistic”; “negative, he’s always causing trouble”). To further understand the difference between these two characters, we applied the text-analysis software package LIWC-22 (Boyd et al., 2022) to participants’ open answers regarding the characters. We then used dependent t-tests to assess the difference in the representation of each of the 9 categories between the two characters. This analysis revealed a significant difference between Clara (M = 2.9, SD = 5.47) and Alexander (M = 0.67, SD = 2.34); t(49) = -2.66, p = 0.01 in the affiliation category, a significant difference between Clara (M = 11.56, SD = 18.69) and Alexander (M = 26.83, SD = 32.23); t(49) = 2.81, p = 0.007 in the affect category; and a significant difference between Clara (M = 7.49, SD = 16.88) and Alexander (M = 24.66, SD = 32.74); t(49) = 3.31, p = 0.0017 in the negative tone category. (Fig. 3C).

### 4.2. Regions Involved in Processing the Narrative

In order to identify brain regions that were involved in processing the narrative, we implemented the ISC analysis (Hasson et al., 2004) on the pre-registered ROIs, as well as on the whole-brain parcellation (122 parcels). We found that listening to the narrative induced neural synchronization in the pre-registered ROIs within the DMN (including left TPJ and PCC, Fig. 4A) as well as in 56 (out of the 122) parcels, including the DMN (PCC, dmPFC, bilateral TPJ and bilateral temporal pole), bilateral auditory cortex, bilateral para-hippocampal gyrus, language-related areas in the ventrolateral prefrontal cortex (vlPFC) and bilateral superior temporal gyri (STG; Fig. 4B).

**Fig. 4.**
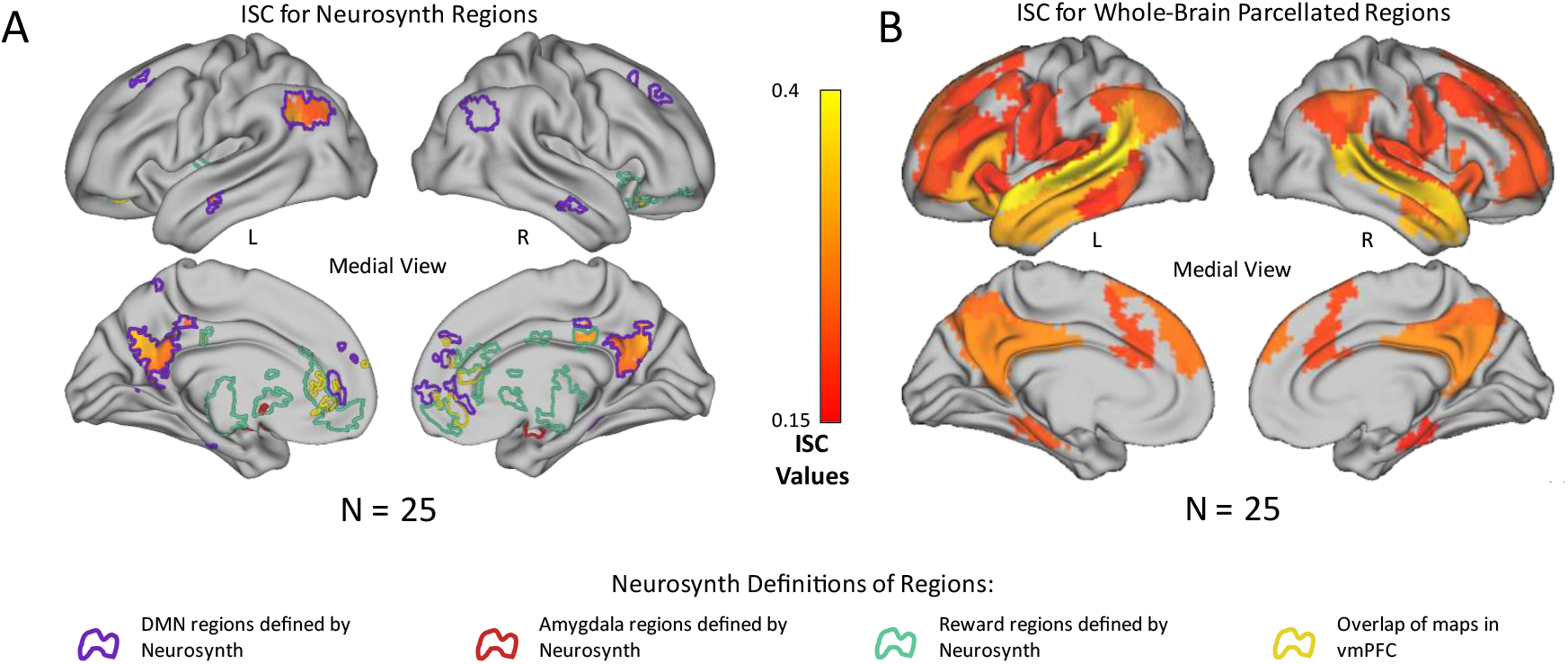
Inter-subject correlation (ISC) values among all participants when listening to the story. A) ISC values within regions defined by Neurosynth. B) ISC values within whole-brain parcels.

### 4.3. Engagement-Dependent Brain Response Synchronization

We used IS-RSA to test whether higher narrative engagement was associated with more similar neural responses. This analysis revealed that in the pre-registered ROIs, the more the participants were engaged, the more synchronized their neural response was in the DMN (including the left TPJ and the PCC) and in reward areas (including the anterior cingulate, Fig. 5A). Moreover, exploratory whole-brain IS-RSA analysis revealed that increased engagement was associated with increased synchronization in 29 parcels, including bilateral TPJ, right STG, dmPFC and left dlPFC (Fig. 5C). Of these parcels, 9 were associated with the control network, 6 with the DMN, 5 with the dorsal attention network (DAN), 4 with the ventral attention network (VAN), 2 with the limbic network, and 1 with each of the somatomotor, temporal parietal and visual networks (see supplemental Table 1). Figs. 5B and 5D display the overlap of parcels involved in processing the narrative among the Low-Engaged, Medium-Engaged and Highly Engaged groups (i.e., the parcels with ISC values that exceeded the threshold within at least one of these groups). For example, whereas the right superior temporal gyrus (STG) was involved in processing the narrative in all groups (Fig. 5D, marked in red, and Fig. 5E), the ventral posterior cingulate cortex (PCC) was involved in processing the narrative only in the medium- and highly engaged groups (but not the low-engaged group; Fig. 5D, marked in yellow, and Fig. 5E); and the left dorsolateral prefrontal cortex (dlPFC) was involved in processing the narrative only in the highly-engaged participants (Fig. 5D, marked in green, and Fig. 5E).

**Fig. 5.**
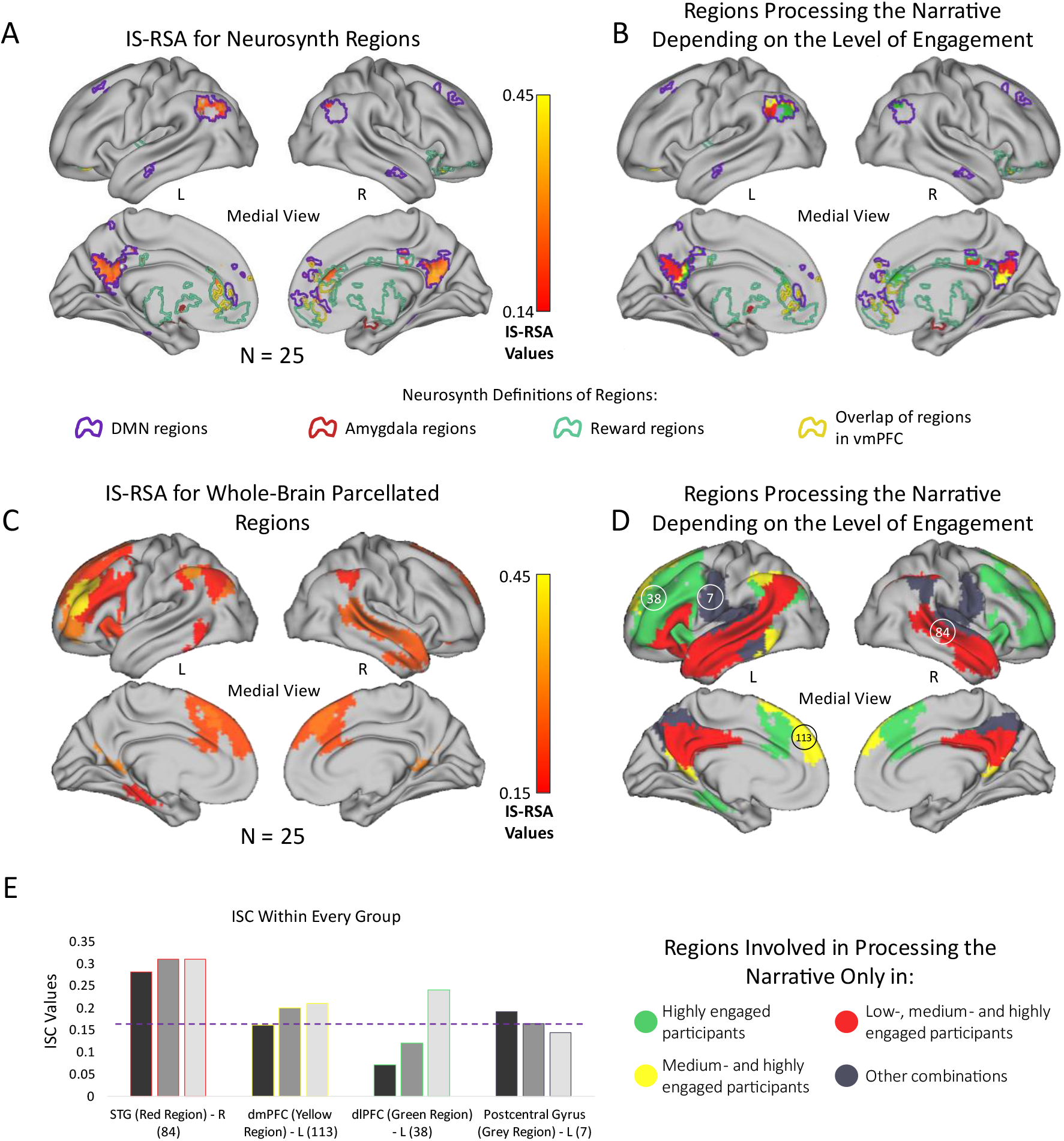
Engagement-dependent brain response synchronization. A) Inter-Subject Representational Similarity Analysis (IS-RSA) in regions defined by Neurosynth. B) Overlap of Neurosynth regions that passed a threshold (the 99 percentile of the null-distribution generated for ISC values) within every subgroup (low-engaged participants, medium-engaged participants, and highly engaged participants). C) IS-RSA values in whole-brain parcellation. D) Overlap of parcels that passed the threshold (99 percentile of the null-distribution generated for ISC values) within every subgroup. Parcels 7, 38, 84 and 88 represent the Postcentral Gyrus, the dorsolateral Prefrontal Cortex (dlPFC), the Superior Temporal Gyrus and the Posterior Cingulate Cortex, respectively. E) The ISC values of every subgroup within four selected parcels displayed on a bar plot, demonstrating the four possibilities for overlaps. The dashed line represents the threshold.

### 4.4. Valence-Engagement-Dependent Brain Response Synchronization

To test the influence of engagement valence on the neural responses to the narrative, we applied IS-RSA to timepoints in which a character inducing positive engagement appeared (Clara) and separately on timepoints in which a character inducing negative engagement appeared (Alexander). We found that increased positive engagement was associated with increased synchronization in many brain regions including left TPJ, medial PFC and posterior cingulate (Fig. 6A and 6B). On the other hand, increased negative engagement was associated with increased synchronization in the bilateral anterior insula, regions in the somatosensory cortex and right dlPFC (Fig. 6B). Both positive and negative engagement were associated with increased synchronization in the left dlPFC (Fig. 6B).

**Fig. 6.**
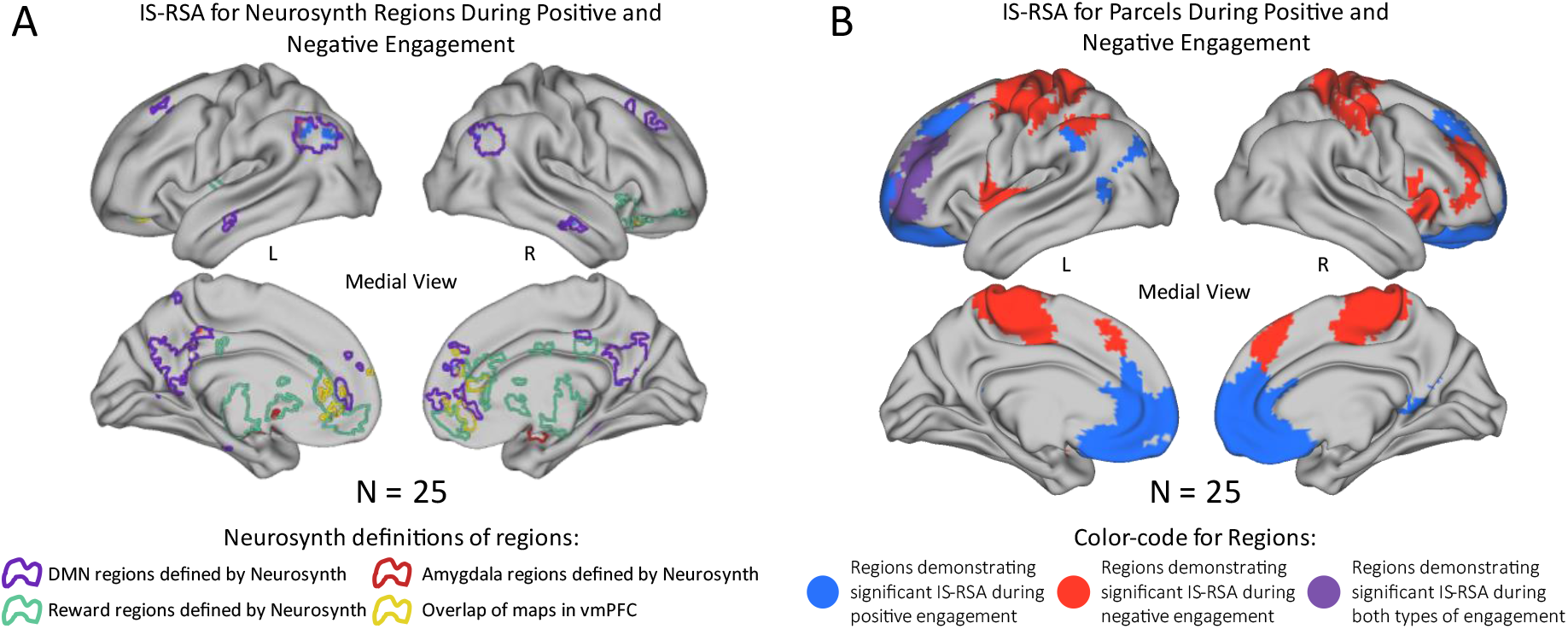
Character-engagement-dependent brain response synchronization. A) Inter-Subject Representational Similarity Analysis (IS-RSA) in regions defined by Neurosynth during Clara’s (blue), Alexander’s (red) or both (purple) appearances. B) Inter-Subject Representational Similarity Analysis (IS-RSA) in whole-brain parcellation during Clara’s (blue), Alexander’s (red) or both (purple) appearances.

### 4.5. Engagement-Dependent Functional Connectivity

We tested whether higher narrative engagement was associated with stronger connectivity between regions that demonstrated engagement-dependent response. To do that, we combined the ISFC analysis (Simony et al., 2016) with RSA analysis (Finn et al., 2020). The resulting ISFC-RSA analysis demonstrated a significant correlation between narrative engagement and functional connectivity in 4 edges between the pre-registered ROIs: 3 connected between regions of the DMN and 1 connected between reward regions and the vmPFC (Fig. 7A). Moreover, in the whole-brain parcellation, this analysis revealed 22 edges: 2 connected nodes within the DMN, 5 connected nodes within the DAN, 5 within the CN and 1 within the limbic network (Fig. 7C). Networks that were connected shared a single edge, except for the DMN and the control network, which shared 4 edges (Fig. 7A).

**Fig. 7.**
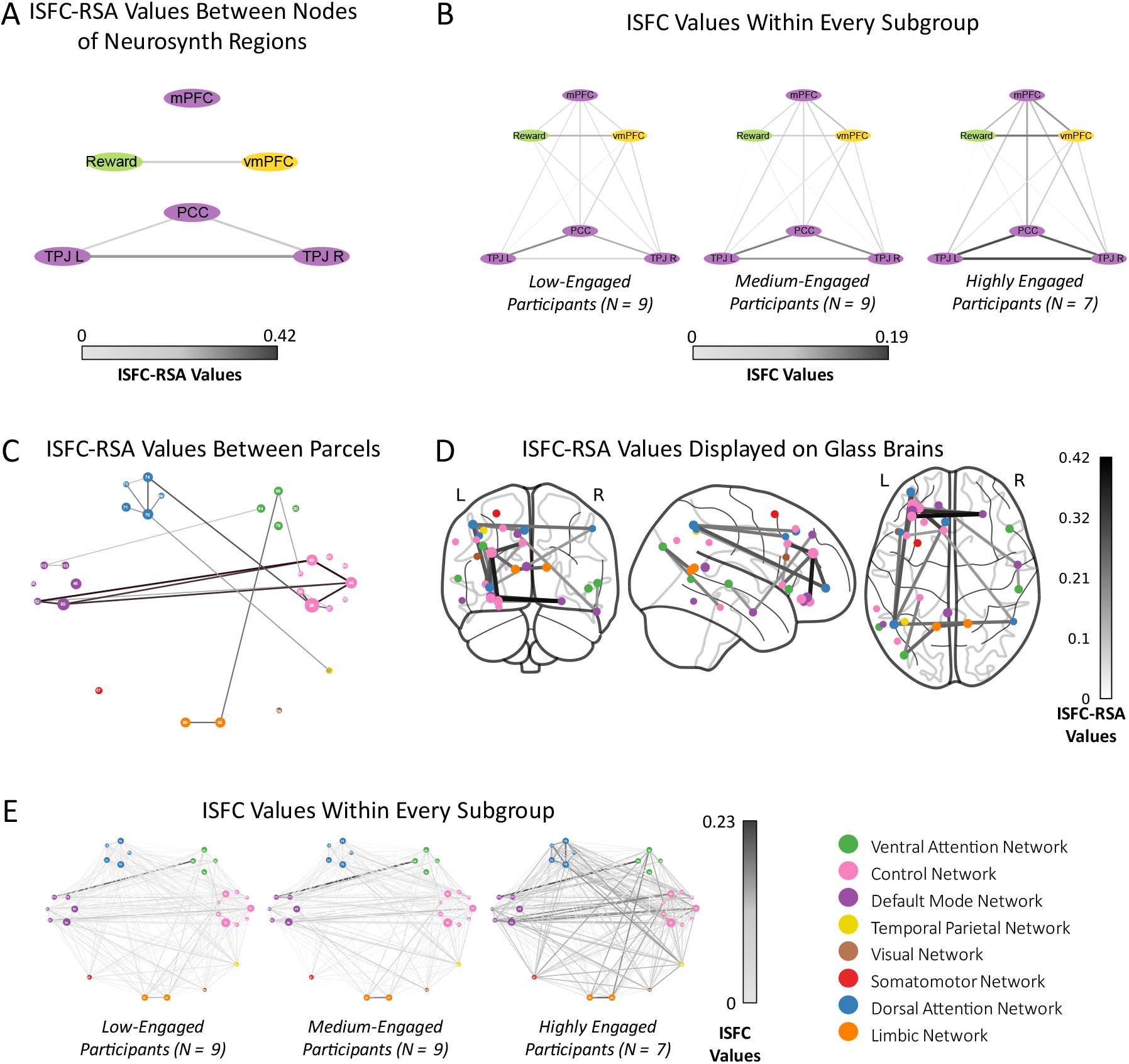
Inter-subject functional connectivity combined with representational similarity analysis. A) ISFC-RSA results displayed on a graph. Edges’ width and color represent the extent of the engagement dependency. Nodes are Neurosynth regions that have been found significant in the IS-RSA analysis (Fig. 5). B) ISFC values within every subgroup (low-engaged participants, medium-engaged participants, and highly engaged participants). C) ISFC-RSA results displayed on a graph. Edges’ width and color represent the extent of the engagement dependency. Nodes are parcels that have been found significant in the IS-RSA analysis (Fig. 5), their diameter corresponding to the IS-RSA values, their color indicating their association with the networks. D) ISFC-RSA results displayed on glass brains. E) ISFC values within every subgroup (low-engaged participants, medium-engaged participants, and highly engaged participants).

## 5. Discussion

A certain narrative may immerse one individual in the plot, while another may feel bored and despondent. In the present study we asked how engagement with the narrative shapes individuals’ brain response. We found that narrative engagement synchronized listeners’ brain responses in many brain regions, including parts of the DMN, dlPFC and the reward system. Moreover, positive engagement was associated with higher synchronization within medial prefrontal regions, whereas negative engagement was associated with higher synchronization in the anterior insula and somatosensory regions. Finally, being engaged with a narrative strengthened the connectivity within and between the DMN and the control and dorsal attention networks.

The DMN is known to be involved in narrative comprehension and interpretation (Deniz et al., 2019; Nguyen et al., 2019; Simony et al., 2016; Yeshurun et al., 2021). Consistent with our hypothesis, we found that the more individuals were engaged with the narrative, the more synchronized was their response in regions within the DMN, including dmPFC, bilateral TPJ and right temporal pole (Fig. 5). This is in line with previous studies that found the response within these regions to be more synchronized for more (vs. less) engaging narratives (Cohen et al., 2017; Grall et al., 2021; Imhof et al., 2017; Schmälzle et al., 2015) and for the engaging parts of a specific narrative (Nummenmaa et al., 2012, 2014; Song et al., 2021). Our study further suggests that these DMN regions’ synchronization reflects individual differences in engagement, and not merely group-level states of engagement. Interestingly, our results also revealed that engagement synchronized the response in regions outside the DMN, such as the left dlPFC, the left para-hippocampal gyrus, and the right STG (Fig. 5C). Previous studies suggested that these regions are involved in verbal working memory and narrative comprehension (Coelho et al., 2012; Hartley & Speer, 2000; Klaus & Schutter, 2018; Meyer et al., 2012; Regev et al., 2013; Shin et al., 2015), as well as in episodic memory and recollection of events (Diana et al., 2010; Dickerson & Eichenbaum, 2010; Hasson et al., 2008; Ranganath & Ritchey, 2012; Song et al., 2021; Staresina et al., 2011). Thus, our results suggest that engagement synchronizes the response of regions involved in reconstruction and integration of the narrative.

There was a dissociation in engagement-dependent neural synchronization during appearances of a character that induced positive engagement (Clara) versus a character that induced negative engagement (Alexander). This difference might stem from the contrast in identification with the characters and the valence of emotions the characters elicited (see Figs. 3B and 3C). We found that positive engagement increased synchronization in regions within the DMN, especially the vmPFC, and with ACC reward regions (Fig. 6). The neural activity of the DMN has been shown to be influenced by identification with a story’s characters (Broom et al., 2021), and hence the engagement-dependent synchronization observed during Clara’s appearance may have been mediated by participants’ identification with her. Moreover, the vmPFC has been associated with many functions, including reward processing (Cauda et al., 2011; Pujara et al., 2016), social cognition (Molenberghs et al., 2016; Shamay-Tsoory et al., 2009) and interpretation of socio-emotional qualities of narratives (Burin et al., 2014; for a review see Hiser & Koenigs, 2018). Our findings of engagement-dependent synchronization of reward regions (Fig. 6A) suggest an engagement-dependent reward effect, which is consistent with behavioral studies (Green et al., 2004). Moreover, our findings of engagement-dependent connectivity between these reward regions and the vmPFC (Fig. 7A) may propose a mechanism for the association between narrative understanding and the enjoyment it produces among engaged participants.

Whereas positive engagement was associated with neural synchronization within DMN and reward regions, negative engagement was associated with increased neural synchronization in the insula and somatosensory regions. Participants’ reactions to Alexander, the character that elicited negative responses, included more affect, more negative tone and less affiliation than participants’ reactions to Clara. Previous literature suggests that processing stimuli that elicit immorality, disgust and other aversive emotions activates the insula (Chapman & Anderson, 2013; Chapman et al., 2009; Denke et al., 2014; Jones & Fitness, 2008; Sanfey et al., 2003; Wicker et al., 2003, Wright et al., 2004; Young & Saxe, 2011) and somatosensory cortices (Adolphs, 2002; Koenigs et al., 2007; Kropf et al., 2018; Straube & Miltner, 2011). Thus, our findings imply that engagement during the appearance of a negative character synchronized listeners’ neural response in regions involved in negative reaction to stimuli.

Engagement with the narrative increased functional connectivity within and between different networks: DMN, DAN and the CN (Fig. 7C). Functional connectivity within regions of the DMN has been shown to be enhanced when a story was more comprehensible (Simony et al., 2016), and this connectivity has been proposed to be influenced by context (Yeshurun et al., 2021). Our findings further indicate that narrative engagement synchronizes the response of the different regions of the DMN, regardless of narrative comprehensibility. The DAN has been associated with both bottom-up and top-down processes of involuntary and voluntary attention, respectively (Bowling et al., 2020). We suggest that enhanced connectivity within this network may have had a role in eliciting the attentional-focus aspect of engagement (Busselle & Bilandzic, 2009). This attentional focus could have been both involuntary (Ki et al., 2016) and voluntary (Regev et al., 2019), demonstrating a complex processing of the stimulus. The CN is thought to adjust other neural networks, as it is functionally connected with them (Marek & Dosenbach, 2018). Specifically, throughout different types of tasks, the CN’s connectivity pattern shifts between different neural networks (Cole et al., 2013). Thus, the enhanced connectivity within the CN in the present study may imply that it integrated information coming from other networks, and this is in line with the finding of correlation between suspense ratings during a story and synchronization within the CN during the story (Schmälzle & Grall, 2020).

Our findings revealed engagement-dependent functional connectivity, not just within brain networks, but also between the DMN, CN and DAN (Fig. 7C). These networks have been traditionally referred to as “task-negative” (DMN; Buckner, 2012; Buckner et al., 1995; Raichle et al., 2001) and “task-positive” (CN and DAN; Fox et al., 2005; Knyazev et al., 2016). The “task-negative” network was shown to be non-synchronized with the “task-positive” networks during rest (Fox et al., 2009; Fox & Raichle, 2007). Our finding that engagement with the narrative increased the functional connectivity between these networks aligns with other findings (Aboud et al., 2019; Bailey et al., 2018) and suggests that these networks are coupled when processing naturalistic stimuli. Moreover, the enhanced connectivity between the CN and DMN and DAN may be inferred as mediating information between the DMN and DAN, and consequently mediating between attention and comprehension of the story.

## 6. Conclusion

Taken together, our findings reveal that engagement with a narrative influences processes of mentalizing, reward, working memory and attention. Focusing on individual differences in engagement ratings allowed us to offer a complementary perspective to existing literature, showing that these processes have been detected due to engagement per se, and not due to differences in the narrative’s content. These findings pave the path for future research, which could examine engagement in more interactive settings, as well as individual differences of engagement tendencies and their prediction of psychological health and emotion-regulation strategies.

## 7. Declaration of Competing Interest

The authors state that they have no competing financial interests or personal relationships that could have affected the work disclosed in this study.

## 8. Authors Contributions

Tal Ohad: Conceptualization, Methodology, Software, Writing – original draft; Yaara Yeshurun: Conceptualization, Methodology, Software, Writing – original draft, review, editing, Supervision.

## Acknowledgements

We would like to thank Claire Chang and Christina Lazaridi for fruitful discussions along the way.

## Funding

This research was supported by the Israel Science Foundation, grant No. 2434/19 (YY).

## 9. Supplementary

**Table 1.**
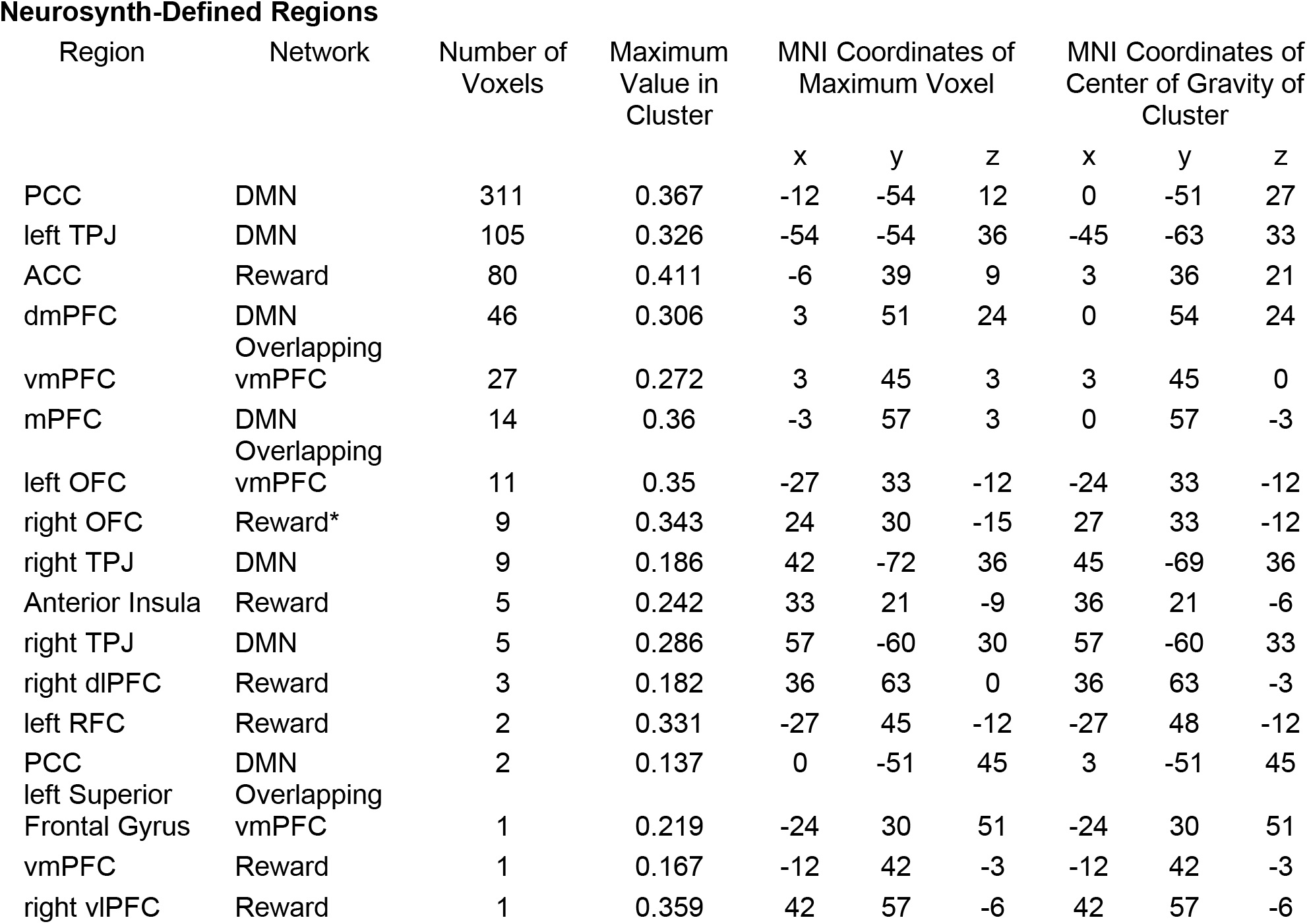
Clusters of voxels with an engagement-dependent response. For each cluster, the table indicates the region in the brain, network it is associated with, number of its voxels, maximal IS-RSA value, coordinates of the voxel carrying the maximal IS-RSA value and coordinates of the center of gravity of the cluster.

**Table 2.**
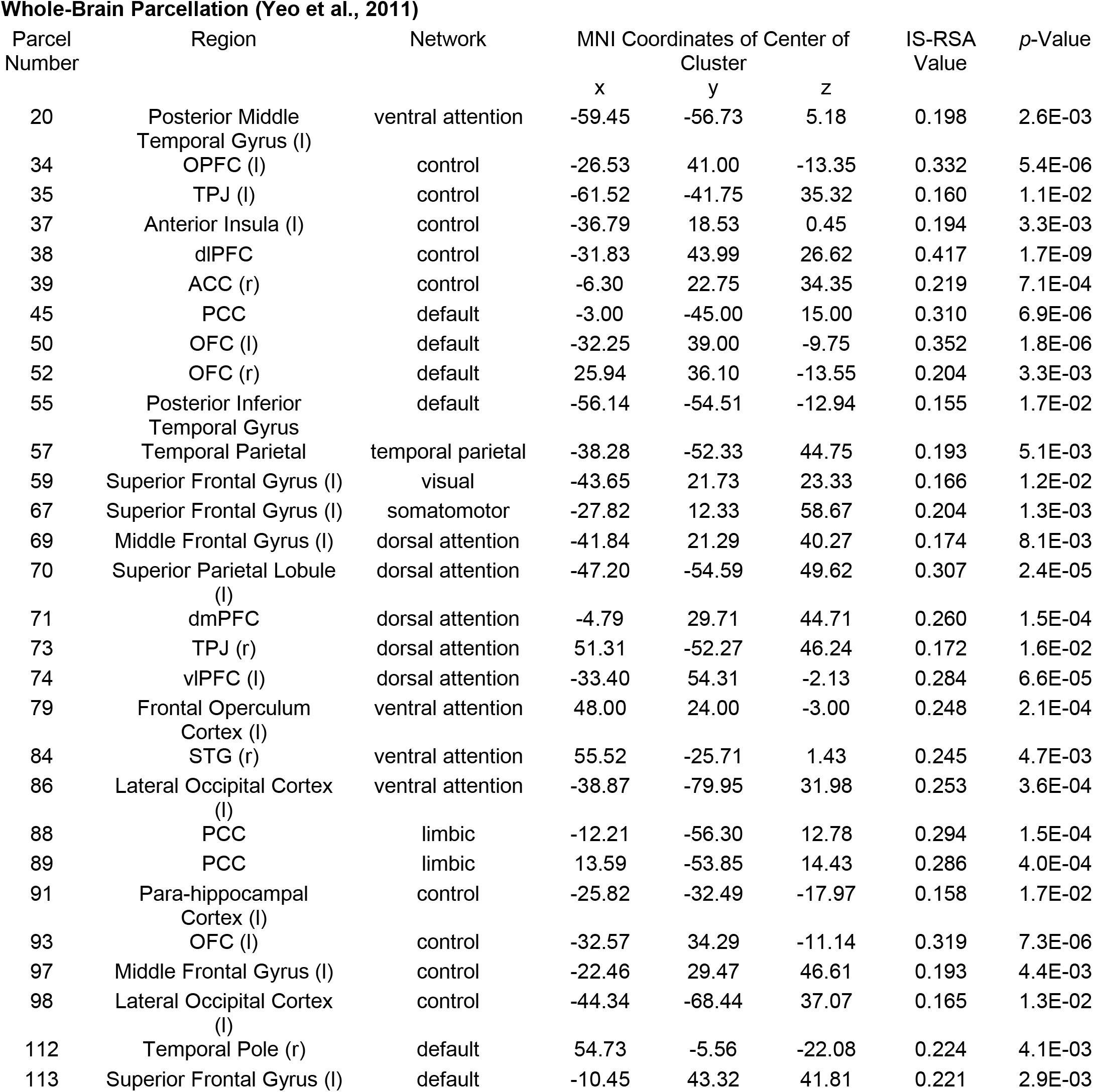
Parcels with a significant engagement-dependent response. For each parcel, the table indicates its conventional number, region in the brain, network it is associated with, MNI coordinates of the center of the parcel, IS-RSA value, and *p*-values of the IS-RSA values.

| Parcel Number | Region | Network | MNI Coordinates of Center of Cluster |  |  | IS-RSA Value | *p*-Value |
| --- | --- | --- | --- | --- | --- | --- | --- |
|  |  |  | x | y | z |  |  |
| 20 | Posterior Middle Temporal Gyrus (l) | ventral attention | -59.45 | -56.73 | 5.18 | 0.198 | 2.6E-03 |
| 34 | OPFC (l) | control | -26.53 | 41.00 | -13.35 | 0.332 | 5.4E-06 |
| 35 | TPJ (l) | control | -61.52 | -41.75 | 35.32 | 0.160 | 1.1E-02 |
| 37 | Anterior Insula (l) | control | -36.79 | 18.53 | 0.45 | 0.194 | 3.3E-03 |
| 38 | dIPFC | control | -31.83 | 43.99 | 26.62 | 0.417 | 1.7E-09 |
| 39 | ACC (r) | control | -6.30 | 22.75 | 34.35 | 0.219 | 7.1E-04 |
| 45 | PCC | default | -3.00 | -45.00 | 15.00 | 0.310 | 6.9E-06 |
| 50 | OFC (l) | default | -32.25 | 39.00 | -9.75 | 0.352 | 1.8E-06 |
| 52 | OFC (r) | default | 25.94 | 36.10 | -13.55 | 0.204 | 3.3E-03 |
| 55 | Posterior Inferior Temporal Gyrus | default | -56.14 | -54.51 | -12.94 | 0.155 | 1.7E-02 |
| 57 | Temporal Parietal | temporal parietal | -38.28 | -52.33 | 44.75 | 0.193 | 5.1E-03 |
| 59 | Superior Frontal Gyrus (l) | visual | -43.65 | 21.73 | 23.33 | 0.166 | 1.2E-02 |
| 67 | Superior Frontal Gyrus (l) | somatomotor | -27.82 | 12.33 | 58.67 | 0.204 | 1.3E-03 |
| 69 | Middle Frontal Gyrus (l) | dorsal attention | -41.84 | 21.29 | 40.27 | 0.174 | 8.1E-03 |
| 70 | Superior Parietal Lobule (l) | dorsal attention | -47.20 | -54.59 | 49.62 | 0.307 | 2.4E-05 |
| 71 | dmPFC | dorsal attention | -4.79 | 29.71 | 44.71 | 0.260 | 1.5E-04 |
| 73 | TPJ (r) | dorsal attention | 51.31 | -52.27 | 46.24 | 0.172 | 1.6E-02 |
| 74 | vIPFC (l) | dorsal attention | -33.40 | 54.31 | -2.13 | 0.284 | 6.6E-05 |
| 79 | Frontal Operculum Cortex (l) | ventral attention | 48.00 | 24.00 | -3.00 | 0.248 | 2.1E-04 |
| 84 | STG (r) | ventral attention | 55.52 | -25.71 | 1.43 | 0.245 | 4.7E-03 |
| 86 | Lateral Occipital Cortex (l) | ventral attention | -38.87 | -79.95 | 31.98 | 0.253 | 3.6E-04 |
| 88 | PCC | limbic | -12.21 | -56.30 | 12.78 | 0.294 | 1.5E-04 |
| 89 | PCC | limbic | 13.59 | -53.85 | 14.43 | 0.286 | 4.0E-04 |
| 91 | Para-hippocampal Cortex (l) | control | -25.82 | -32.49 | -17.97 | 0.158 | 1.7E-02 |
| 93 | OFC (l) | control | -32.57 | 34.29 | -11.14 | 0.319 | 7.3E-06 |
| 97 | Middle Frontal Gyrus (l) | control | -22.46 | 29.47 | 46.61 | 0.193 | 4.4E-03 |
| 98 | Lateral Occipital Cortex (l) | control | -44.34 | -68.44 | 37.07 | 0.165 | 1.3E-02 |
| 112 | Temporal Pole (r) | default | 54.73 | -5.56 | -22.08 | 0.224 | 4.1E-03 |
| 113 | Superior Frontal Gyrus (l) | default | -10.45 | 43.32 | 41.81 | 0.221 | 2.9E-03 |

## Notes

### Competing Interest Statement

The authors have declared no competing interest.

